# InterPET: A Curated Benchmark of Sequence Embeddings and Graph Architectures with Interpretability and Biological Validation for PETase Activity Prediction

**DOI:** 10.64898/2026.08.18.745360

**Authors:** Carissa Handrian, Indira Prakoso

**Affiliations:** Independent Researcher, Jakarta, Indonesia; Bioinformatics Research Center, Indonesian Institute of Bioinformatics (INBIO Indonesia), Malang, East Java, Indonesia

**Keywords:** Bioinformatics, machine learning, PETase, protein function prediction, protein sequence analysis

## Abstract

**Motivation:** Machine learning has emerged as a powerful accelerator for identifying PET-hydrolyzing enzymes (PETases). Yet, published models are often evaluated on benchmark performance alone, leaving their biological validity unexamined. Here we present InterPET, a curated benchmark and ablation study addressing both issues.

**Results:** We aggregated sequences from four datasets (PlasticDB, PAZy, PlasticEnz, PEZY-miner), removing duplicate sequences, and filter data leakage, yielding a training set of 937 sequences and a benchmark of 139 sequences. Eight model configurations were trained and evaluated, spanning three embeddings (ESM-2, ProtT5, classical AAC/CTD descriptors), two tree-based classifiers (XGBoost, Random Forest), and two GraphSAGE variants differing in sequence-only and sequene plus 3D structure data. ESM-2 + XGBoost achieved the best performance (F1 = 0.91, AUC = 0.99, MCC = 0.90). SHAP-based feature attribution linked top-ranked AAC/CTD features (proline content, solvent accessibility, hydrophobicity) to known determinants of PETase activity, and cross-representation correlation showed that embedding-based models implicitly re-encode much of the same biophysical signal. However, in-silico mutagenesis revealed that the top-ranked M1 recovered only 0.5/3 catalytic-triad residues. These findings demonstrate that representation choice, classifier architecture, and evaluation criteria interact in ways a single leaderboard metric cannot capture.

**Availability and implementation:** InterPET datasets and code are available at https://github.com/indi-raprakoso/interpet/.

## 1 Introduction

One of the most prominent plastics currently is polyethylene terephthalate (PET), a type of polyester plastic polymerized from ethylene glycol (EG) and terephthalic acid (TPA). To address the problem of PET pollution, ongoing research is focused on PET degradation into smaller, recyclable products. Chemical degradation using alcoholysis holds potential in PET degradation, Yet, it is costly and technically challenging. PET is relatively resistant to mechanical stress and heat due to its polymer strength, making these two degradation methods also ineffective (Jiao *et al*. 2024). Recently, the microorganism *Ideonella sakaiensis* was identified to be an organism that uses PET as its main energy and carbon source (Yoshida *et al*. 2021). This opened up the future path for sustainable PET degradation.

However, the vast sequence space of potential enzymes renders traditional, wet-lab screening methods prohibitively time-consuming and expensive. In this context, computational biology and machine learning (ML) have emerged as powerful accelerators, enabling the rapid in silico prediction and prioritization of promising enzyme candidates (Lu *et al*. 2022). Despite their immense potential, the application of ML to PETase discovery is fraught with challenges that are often overlooked. The first critical hurdle is the quality and integrity of the data itself. These include: Positive-Unlabeled (PU) Contamination, where the absence of an enzyme in a “plastic-active” database is incorrectly treated as a negative label, despite the lack of experimental evidence confirming its inactivity (Claesen *et al*. 2015); (2) Data Leakage, where sequence or structural similarities between training and test sets artificially inflate performance metrics (Sasse *et al*. 2025); and (3) Artifact Anisotropy, the presence of non-biological, systematic biases in the data (Schnurr *et al*. 2019). Addressing these biases through rigorous, explicit curation is therefore the first pre-requisite for any trustworthy PETase predictor.

Furthermore, even with clean data, the optimal representation and modeling strategy for enzymatic function remains an open question. While powerful protein Language Models (pLMs) like ESM-2 and ProtT5 have revolutionized the field by generating rich, contextualized embeddings from sequence alone, they operate as “black boxes”. Conversely, classical, hand-crafted features like Amino Acid Composition (AAC) and Composition-Transition-Distribution (CTD) are more interpretable but may lack the nuanced information captured by pLMs (Fan *et al*. 2026). Additionally, although the integration of three-dimensional contact maps could potentially improve predictive performance, this avenue remains underexplored. A systematic comparison of various embedding types, classifiers, and graph topologies within a unified benchmark could therefore shed light on a question that has yet to be conclusively addressed.

Crucially, the interpretability must be checked against known biology, since a model can achieve strong benchmark accuracy while relying on signal unrelated to the experimentally confirmed catalytic machinery. Based on the background above, this research aims to develop and systematically benchmark machine learning models for predicting PET-hydrolase (PETase) activity from sequence. We present InterPET, a benchmarking framework built on three complementary contributions that mirror the challenges outlined above (Figure 1).

**Figure 1.**
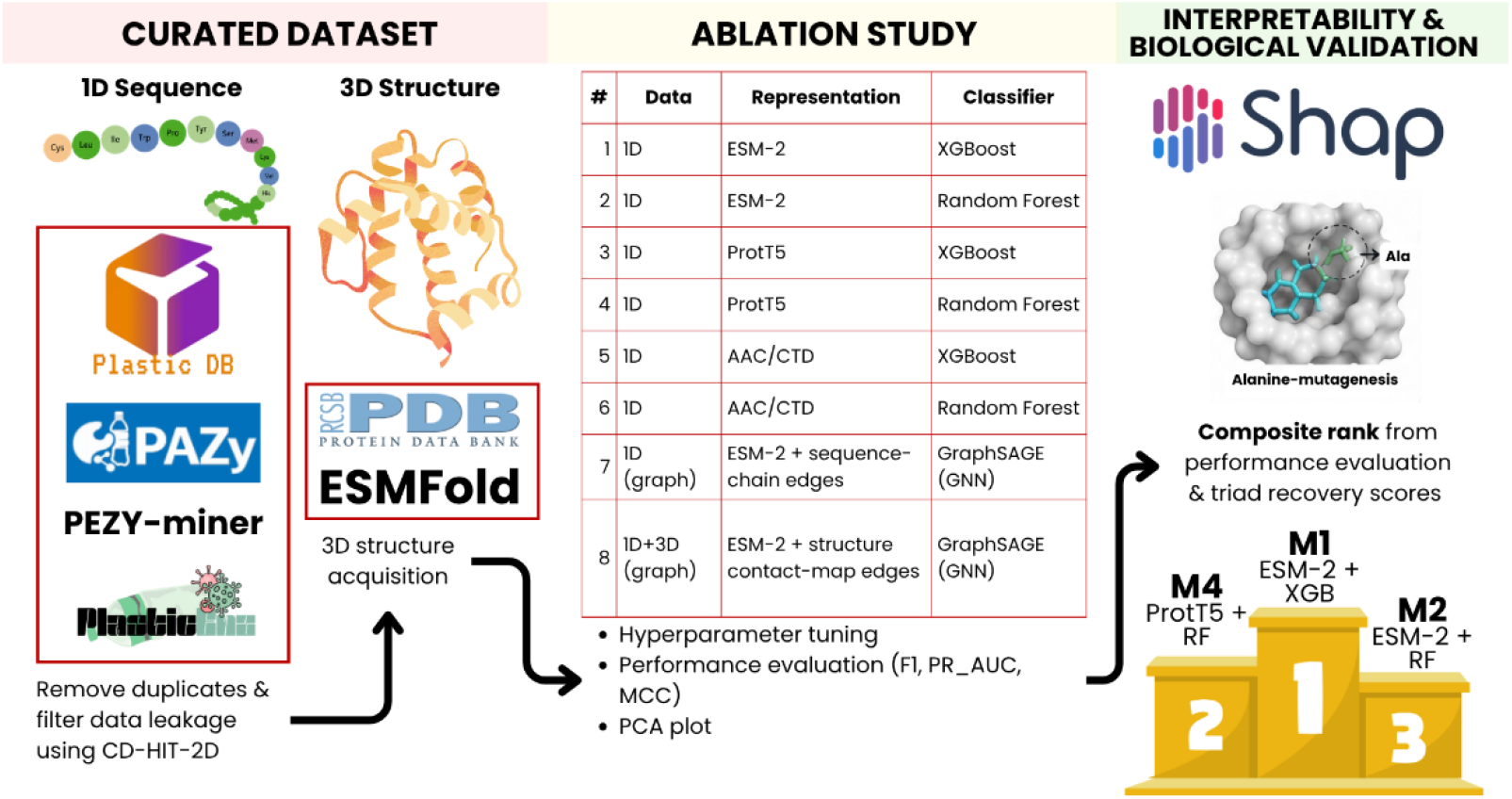
Graphical overview of the InterPET workflow. The pipeline consists of three stages. (Left) Curated dataset: protein sequences (1D) and structures (3D). (Center) Ablation study: eight model configurations (M1–M8) are trained and evaluated. All models undergo hyperparameter tuning, performance evaluation (F1, PR-AUC, MCC), and PCA-based representation analysis. (Right) Interpretability and biological validation: SHAP-based feature attribution and in-silico alanine-scanning mutagenesis. Models are ranked by a composite score combining benchmark performance and catalytic-triad recovery, with ESM-2 + XGBoost (M1), ESM-2 + Random Forest (M2), and ProtT5 + Random Forest (M4) among the top-performing configurations.

## 2 Methods

### 2.1 Dataset Curation and Benchmark Construction

The sequence data were aggregated from four datasets, including PlasticDB (Gambarini *et al*. 2022), PAZy (Buchholz *et al*. 2022), PlasticEnz (Krzynowek, Snoeks, and Faust 2026), and PEZY-miner (Jiang *et al*. 2023) to construct the training set. Negative dataset was defined as enzymes experimentally documented as active on non-PET plastic substrates rather than randomly sampled non-enzymatic sequences. The PlasticEnz test split was reserved as an independent benchmark and kept entirely separate throughout curation.

All sequences shorter than 20 amino acids were discarded. Sequences with contradictory labels across sources were excluded, and exact duplicates were removed. CD-HIT-2D was applied to remove negative sequences ≥90% identical to any positive sequence, and a second CD-HIT-2D run filtered out training sequences ≥90% similar to the benchmark (Li and Godzik 2006, Bursteinas *et al*. 2016). Finally, ESM-2 embeddings were computed for all sequences, corrected for anisotropy by subtracting the global mean vector, and visualized via UMAP to assess class separation (McInnes *et al*. 2018, Lin *et al*. 2023).

### 2.2 3D Structure Acquisition

For each sequence, an initial search was performed using the RCSB Sequence Search API (Piehl *et al*. 2025) with a 95% identity cutoff and an E-value cutoff of 1.0. For sequences lacking a sufficiently identical experimental structure, ESMFold (facebook/esmfold_v1) was employed as a prediction-based fallback, with sequences exceeding 500 residues truncated prior to prediction and mean pLDDT scores computed as confidence metrics (Lin *et al*. 2023).

### 2.3 Ablation Study Design

The ablation of eight model configurations were designed to disentangle the contribution of sequence representation, classifier architecture, and structural information to PETase prediction (Table 1). Three sequence representation types were compared: ESM-2 (Lin *et al*. 2023) and ProtT5 (Elnaggar *et al*. 2022), two pretrained protein language models differing in architecture and pretraining corpus, and AAC/CTD (Dubchak *et al*. 1995, Malik *et al*. 2022), a classical handcrafted descriptor based on amino-acid composition and transition/distribution statistics. This comparison isolates whether a learned deep representation is necessary or whether simpler and fully interpretable composition features suffice. Each embedding-based representation (ESM-2, ProtT5) was paired with two tree-based classifiers, XGBoost (Chen and Guestrin 2016) and Random Forest (Breiman 2001, Krzynowek, Snoeks, and Faust 2026), yielding four tabular models (#1–4) plus two additional AAC/CTD-based models (#5–6), for a total of six 1D (sequence-only) configurations. A Graph Neural Network (two-layer GraphSAGE) (Hamilton, Ying, and Leskovec 2017) was used to test the contribution of structure directly. Two graph variants were constructed with identical per-residue ESM-2 node features, architecture, and training procedure, differing only in edge topology. Model #7 used sequence-adjacency (“chain”) edges as a structure-free control, while Model #8 used Cα–Cα contact-map edges (<8 Å) derived from the acquired 3D structures (Xu and Bonvin 2024, Nguyen *et al*. 2025). This 1D vs 1D+3D comparison (#7 vs. #8) provides the cleanest possible isolation of whether 3D structural information improves prediction beyond graph architecture alone.

**Table 1.** Ablation Study Models.

| # | Data | Representation | Classifier |
| --- | --- | --- | --- |
| 1 | 1D | ESM-2 | XGBoost |
| 2 | 1D | ESM-2 | Random Forest |
| 3 | 1D | ProtT5 | XGBoost |
| 4 | 1D | ProtT5 | Random Forest |
| 5 | 1D | AAC/CTD | XGBoost |
| 6 | 1D | AAC/CTD | Random Forest |
| 7 | 1D (graph) | ESM-2 + sequence-chain edges | GraphSAGE (GNN) |
| 8 | 1D+3D (graph) | ESM-2 + structure contact-map edges | GraphSAGE (GNN) |

### 2.4 Model Optimization & Evaluation

All eight models were trained on the curated training set and tuned using stratified 5-fold cross-validation (random_state = 42) to preserve class balance across folds. For the six tabular models (#1–6), a small grid search (4–5 candidate configurations per model) was performed over classifier-specific hyperparameters. Mean cross-fold F1 was used as the selection criterion, with identical search grids applied across all three representations (ESM-2, ProtT5, AAC/CTD) to keep the comparison fair. The class imbalance was addressed via scale_pos_weight (XGBoost) and class_weight=“balanced” (Random Forest) (Amiri, Afshari, and Soltani 2025). The two graph-based models (#7–8) were tuned over GraphSAGE architecture under the same cross-validation protocol. The final models were then retrained on the full training set with an extended epoch budget (maximum 100 epochs, patience of 15) using the Adam optimizer (learning rate 1e-3, weight decay 1e-4).

For every model, the classification threshold was selected from out-of-fold (OOF) predictions generated during cross-validation to prevent threshold leakage (Perdana, Hantono, and Ferdiana 2026). Each model’s finalized threshold and hyperparameters (Table S1) were then applied once to the held-out benchmark set to obtain the reported performance metrics, with 95% confidence intervals estimated via bootstrap resampling (n = 2,000). Beyond aggregate metrics, model behavior was inspected visually through ROC and Precision–Recall curves, confusion matrices, and calibration curves. Embedding/feature-space structure was examined via PCA projections to check for class separability and potential distributional artifacts.

### 2.5 SHAP Feature Importance

Feature-level interpretability was assessed using SHAP (SHapley Additive exPlanations) with the TreeExplainer implementation for the XGBoost and Random Forest classifiers across all six tabular models (Lundberg *et al*. 2020). However, ESM-2 and ProtT5 are not inherently interpretable in isolation because embedding dimensions are learned latent directions from self-supervised pretraining rather than engineered biophysical properties. AAC/CTD-based models, by contrast, provided directly interpretable SHAP output, since their input features already carry explicit biophysical meaning.

For the graph-based models (#7–8), SHAP was applied via GradientExplainer to the model’s classifier head (a two-layer feed-forward network operating on the pooled graph embedding). This analysis was framed as a model diagnostic which assessing whether the classifier head relies on a small number of dominant pooled-embedding dimensions or distributes importance broadly. An additivity sanity check (comparing the sum of SHAP values against the model’s actual output difference from baseline) validated attribution reliability for this non-standard architecture.

### 2.6 SHAP-based Cross Representation Validation

Using the same top-SHAP-ranked dimensions described above (the 20 highest mean-absolute-SHAP dimensions per model), we tested whether the embedding-based models capture the same biophysical signal as the explicitly computed AAC/CTD features. Each candidate dimension was correlated (Pearson) against all 167 AAC/CTD features (from Model #5) computed on the same training sequences, yielding 3,340 dimension–feature pairs per model (Eck Van *et al*. 2026).

Raw correlations risk being inflated by confounding through the label itself because both the top-SHAP dimensions and the top AAC/CTD features were selected for their predictiveness of the class label. Therefore, additionally computed a partial correlation was done by regressing the linear effect of the label out of both sides before correlating the residuals and corrected both correlation types for multiple testing using the Benjamini– Hochberg FDR procedure (α = 0.05) (Thissen, Steinberg, and Kuang 2002, Sekaran and Zayed 2026). A dimension–feature pair was classified as a strong match when |r_partial| ≥ 0.5 and FDR q < 0.05.

For Models #7 and #8, graph-level pooled embeddings were only available for the subset of training sequences with a successfully resolved 3D structure (937 of the full training set). Row correspondence with the AAC/CTD features was therefore established via explicit 1D-matching rather than positional correspondence, with correlations computed only on the intersection available in both representations (937/937 sequences matched for both models).

### 2.7 In Silico Mutagenesis and Catalytic-Triad Recovery

The in-silico alanine-scanning mutagenesis was performed to validate model predictions against a biologically grounded and to assess whether the recovered signal is genuinely specific to PET-degrading enzymes rather than a generic serine-hydrolase artifact (Bromberg and Rost 2008, Chandra *et al*. 2014). The scanning was conducted on three fixed reference sequences: IsPETase (*Ideonella sakaiensis* PETase, UniProt A0A0K8P6T7, PDB 5XJH) (Yoshida *et al*. 2016, Joo *et al*. 2018) and LCC (Leaf-branch Compost Cutinase, UniProt G9BY57, PDB 4EB0) (Sulaiman *et al*. 2012, 2014) as independent positive controls. The catalytic triads of those enzymes (Ser160/Asp206/His237 and Ser165/Asp210/His242, respectively) are experimentally confirmed by crystal structure and site-directed mutagenesis. Meanwhile, *Candida antarctica* Lipase B (CalB, UniProt P41365, PDB 1TCA; catalytic triad Ser130/Asp212/His249) was used as a negative control which share the same α/β-hydrolase fold and Ser-Asp-His catalytic mechanism but with no PET-degrading activity (Uppenberg *et al*. 1994). The negative control was used to test whether model-derived importance reflects PET-specific mechanism or merely generic active-site geometry.

For each position in each reference sequence, the residue was substituted with alanine and the resulting change in predicted probability (Δ) yielding a full per-residue importance profile. For the graph-based models (#7–8), each mutation required re-embedding the mutated sequence with ESM-2 and rebuilding the corresponding graph. For Model #8 specifically, this additionally required fetching each reference’s experimental 3D structure (RCSB PDB) and reconstructing Cα–Cα contact-map edges via structure-to-sequence alignment, which were then held fixed across all single-position mutations under a simplifying assumption that point mutations do not substantially perturb global backbone geometry.

Catalytic-triad recovery was quantified as the number of known catalytic residues appearing among each model’s top-15 highest-Δ positions (Jayasekara *et al*. 2023). To test whether the triad residues matter jointly beyond their individual contributions, all three were mutated to alanine simultaneously per reference. The resulting combined delta was compared against the sum of the three individual single-residue deltas (synergy score), with statistical significance assessed via a permutation test against a null distribution built from 200 random position triplets. As secondary validation, a windowed (block) mutagenesis scan (window size = 7 residues) and a native-context masked-token substitution using the model’s own language model <mask> token in place of alanine were additionally performed at the catalytic-triad positions of each reference (Lue and Liau 2023). Model-level biological validity was then summarized as the mean catalytic-triad overlap across the two positive-control references (IsPETase, LCC), reported alongside the corresponding overlap at the CalB negative control.

### 2.8 Composite Rank

A composite score was calculated to integrate benchmark performance with biological validity into a single overall ranking. The rank was calculated for each of the eight models across six criteria: four benchmark metrics (F1, AUC, PR-AUC, MCC) computed on the held-out benchmark set, and two biological-validity metrics derived from the positive-control mutagenesis scan (mean catalytic-triad overlap and mean permutation z-score, each averaged across the IsPETase and LCC reference sequences). For each criterion *c*, models were assigned an ordinal rank *R_c(m)* from 1 (best) to *n* = 8 (worst), using standard competition ranking (ties assigned the minimum rank):

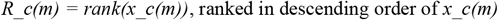

where *x_c(m)* is model *m*’s value for criterion *c*. The composite score, hereafter referred to as the average rank, was then computed as the un-weighted mean of a model’s ranks across all six criteria:

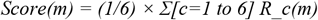

## 3 Results

### 3.1 Dataset Curation

Initial data loading yielded 1,840 raw sequences, comprising 701 positive (PET-degrading) and 1,139 negative (non-PET-degrading) sequences. The distribution across sources was as shown in Table 2. The final training dataset consists of 937 sequences with the ratio 1:1.3 (403:534) between positive and negative dataset. Meanwhile, the final benchmark dataset consists of 139 sequences (22 positive & 117 negative). From 3D acquisition, a total of 1,076 sequences acquire 221 PDB experimental and 855 ESMfold prediction structures. The mean resolution of PDB structures was 1.70 Å, with max 3.17 Å and mean of 99.2% identity. Meanwhile, the mean of ESMfold structures was 86.3 pLDDT with 40 structures lower than 70 pLDDT (min pLDDT = 24.8).

**Table 2.** Distribution of Data Sources.

| Source | Before |  | After |  |
| --- | --- | --- | --- | --- |
|  | Positive | Negative | Positive | Negative |
| PlasticDB | 186 | 143 | 131 | 110 |
| PAZy | 312 | 163 | 239 | 89 |
| PlasticEnz Train | 100 | 506 | 1 | 301 |
| PEZY-miner | 53 | 177 | 32 | 34 |
| **Total Train Set** | **651** | **989** | **403** | **534** |
| PlasticEnz Test (Benchmark) | 50 | 150 | 22 | 117 |
| **Total Dataset** | **701** | **1,139** | **425** | **651** |

Cosine similarity analysis after anisotropy correction revealed near-duplicate patterns only within the training negative set (11 of 534 sequences exceeded the P99 = 1.000 threshold). Meanwhile all other comparisons, including training positive (0/403), training-versus-benchmark (0/937), and within-benchmark (0/139), showed no such cases (Figure S1). The UMAP projection (Figure S2) revealed separated clusters between benchmark and train dataset. No training sequences exceeded the P99 similarity threshold in the train-versus-benchmark comparison, as indicated by the absence of black circles in the UMAP visualization, confirming that the CD-HIT-2D filtering effectively removed near-duplicates and prevented data leakage.

### 3.2 Ablation Model Comparison

Across the eight ablation models evaluated on the curated benchmark set, M1 (ESM-2 + XGBoost) achieved the highest F1 score (0.913), AUC (0.987), and MCC (0.897) among all models (Figure 2), with a 95% bootstrap confidence interval for F1 ranging from 0.810–0.982 and a per-fold cross-validation F1 of 0.935 ± 0.033, indicating stable performance across resampling and cross-validation folds. When grouped by representation family, graph-based models achieved the highest mean benchmark performance (M7-M8; F1 = 0.832, AUC = 0.964), followed by 1D sequence-embedding models (M1–M4; F1 = 0.825, AUC = 0.984) and 1D hand-crafted feature models (M5–M6; F1 = 0.753, AUC = 0.959). Within the tabular models, XGBoost outperformed Random Forest on average across all three feature representations (mean F1 = 0.820 vs. 0.782; mean AUC = 0.978 vs. 0.973), and within the graph family specifically, contact-map edges (Model #8, F1 = 0.840) outperformed sequence-chain edges (Model #7, F1 = 0.824). ROC-AUC values ranged narrowly from 0.951 to 0.987. However, the Precision-Recall curves reveal a substantially wider and more informative spread (AP range: 0.641–0.905). The complete data of benchmark metrics, bootstrap 95% confidence intervals, cross-validation fold metrics, ROC and Precision-Recall curves, confusion matrices, and calibration curves are provided in the Supplementary Table S2-S5 and Supplementary Figure S3-S7.

**Figure 2.**
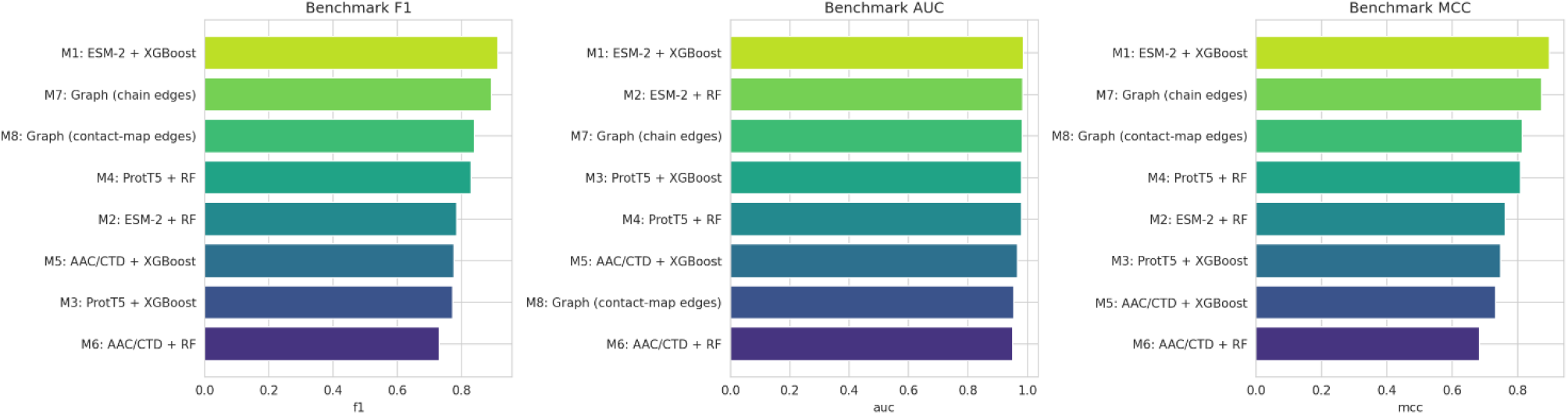
Benchmark comparison of eight model architectures across three evaluation metrics: F1 score (left), area under the ROC curve (AUC, center), and Matthews Correlation Coefficient (MCC, right).

Principal component analysis (PCA) of the input representation of all eight models is shown in Figure S9. In the training-set embeddings of M1–M4 (ESM-2 and ProtT5), the positive and negative classes formed visually distinguishable, partially overlapping clusters along PC1 (18.7–27.7% variance explained), whereas in the corresponding benchmark-set embeddings, the two classes appeared more intermixed with less distinct clustering. For M5–M6 (AAC/CTD features), the two classes largely overlapped in both the training and benchmark sets, with no clearly separated clusters along either PC1 (28.4%) or PC2 (10.6%). For M7–M8 (graph-based pooled embeddings), the training-set points formed two branches diverging from a shared origin, with the positive and negative classes occupying largely separate branches. The benchmark-set embeddings for M7–M8 showed a similar branching pattern, though with fewer points and greater overlap near the branch origin.

### 3.3 Model Interpretability

For the AAC/CTD-based models, whose input features carry explicit biophysical names, the SHAP beeswarm directly identified proline content (AAC_P) as the single most influential feature (mean |SHAP| = 0.894 for M5). The rank followed by solvent-accessibility composition (SolventAccessibility_T12, 0.650), polarity composition (Polarity_T13, 0.507), and secondary-structure composition (SecondaryStructure_C1, 0.337) (Figure 3). Other positively contributing features included AAC_T, AAC_R, Hydrophobicity_C3, and several polarity and polarizability descriptors from the CTD groups.

**Figure 3.**
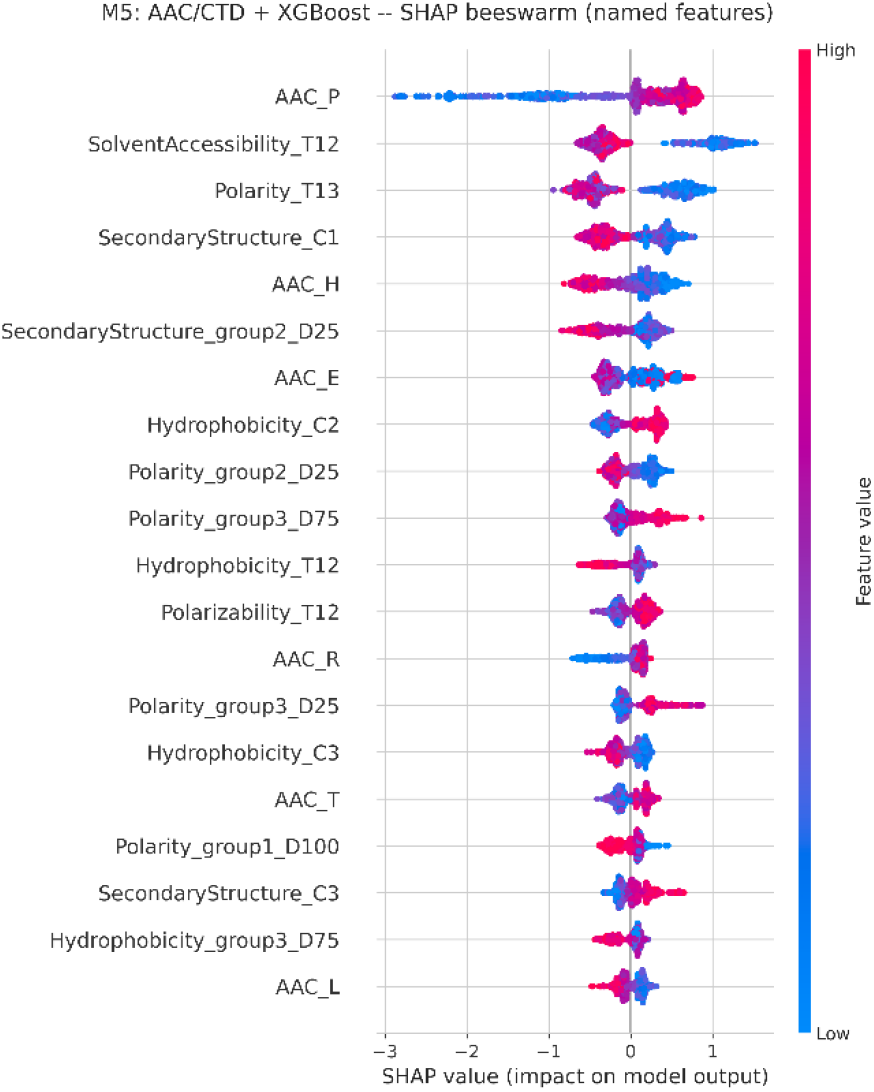
SHAP beeswarm plot for the AAC/CTD. The color indicating the feature’s value (AAC_P = proline composition; SolventAccessibility_T12, Polarity_T13, SecondaryStructure_C1 = composition/transition descriptors) and horizontal position indicating that feature’s SHAP contribution to the predicted probability.

From SHAP-based cross representation validation as shown in Figure S10, across 3,340 dimension–feature pairs tested per model, a substantial fraction remained statistically significant after both FDR correction and label-controlling for ESM-2 (M1: 2,052/3,340 pairs, mean |r| shrinkage = 0.076) and ProtT5 (M3: 1,949/3,340, shrinkage = 0.063), with 40 and 12 pairs respectively meeting a stricter effect-size-and-significance criterion (|r_partial| ≥ 0.5, FDR q < 0.05). In contrast, the graph-based models’ pooled representations showed a markedly different pattern. Despite comparable or higher raw and FDR-significant pair counts (M7: 1,776/3,340; M8: 1,521/3,340), label-controlling produced the largest shrinkage of any model (M7: 0.113; M8: 0.117) and yielded zero pairs meeting the strong-match criterion for either graph variant. This suggests that the apparent correspondence between the GNNs’ pooled graph embeddings and AAC/CTD features is disproportionately attributable to both representations independently tracking the classification label, rather than to a genuine shared biophysical encoding.

### 3.4 Biological Validation

For the IsPETase reference, M3 (ProtT5 + XGBoost) and M4 (ProtT5 + RF) each recovered 1 of 3 catalytic-triad residues within their top-15 ranked positions, with permutation z-scores of 7.2 and 4.3, respectively (p < 0.05), followed by M1 (ESM-2 + XGBoost, z = 3.0, p < 0.05) and M2 (ESM-2 + RF, z = 1.2, not significant) (Figure 4, Table S6). The remaining four models (M5–M8) showed no triad overlap and non-significant z-scores. For the LCC reference, M4 recovered 2 of 3 triad residues (z = 6.8, p < 0.05), while M1, M2, and M3 each recovered 1 of 3 residues, with M3 and M2 also reaching statistical significance (z = 7.6 and 6.7, respectively). Meanwhile, M5–M8 again showed no overlap. For the CalB negative control, M8 (Graph, contact-map edges) showed the highest overlap among all models (2 of 3 CalB catalytic residues in its top-15) with the largest permutation z-score observed across the entire analysis (z ≈ 15.8, p < 0.05), followed by M4, M1, M2, and M7, which also reached statistical significance (z = 2.0–4.5) despite lower overlap counts (0–1 residues). Meanwhile, M3, M5, and M6 showed no significant sensitivity to the CalB reference.

**Figure 4.**
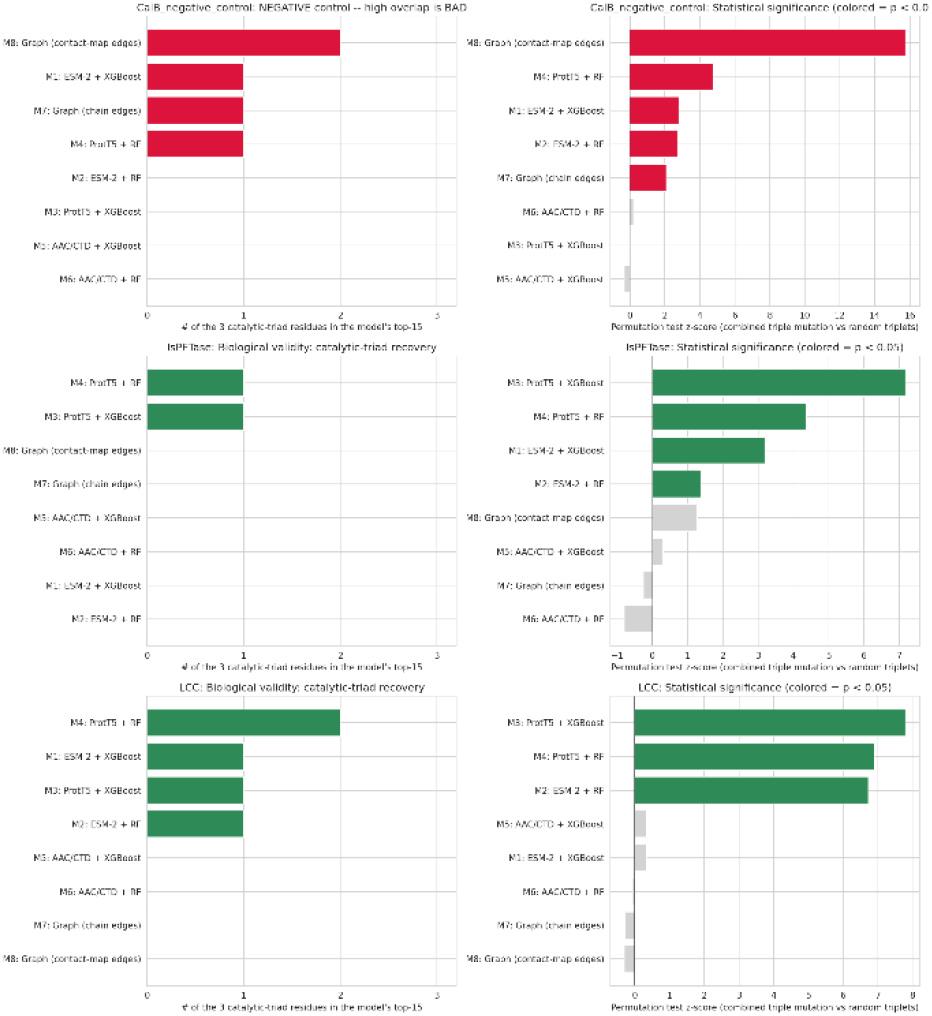
In-silico mutagenesis results across three reference sequences. CalB (negative control; top row), IsPETase (positive control; middle row), and LCC (positive control; bottom row). Left panels = the number of the three catalytic-triad residues recovered within each model’s top-15 most influential positions. Right panels = the permutation test z-score with bars colored to indicate statistical significance (p < 0.05). Green bar indicates desirable, while red bar indicates undesirable off-target sensitivity to a non-PET-active reference’s catalytic residues.

### 3.5 Composite Rank

The composite ranking (Table 3), integrating benchmark performance and biological validity, placed M1 (ESM-2 + XGBoost) first overall (average rank = 1.83), driven by top benchmark scores (F1 = 0.913, AUC = 0.987, MCC = 0.897) despite moderate triad overlap (0.5/3) and z-score (1.76). Meanwhile M4 (ProtT5 + RF) ranked second (average rank = 3.17), excelling in biological validity (triad overlap = 1.5/3, z-score = 5.63) but ranking lower in benchmark metrics. M2 (ESM-2 + RF) and M7 (Graph, chain edges) took third and fourth places, with M7 showing strong predictive accuracy (F1 = 0.894, MCC = 0.875) yet zero triad overlap and a negative z-score (−0.27), indicating its importance scores were no better than random.

**Table 3.** Composite rank of PETase activity prediction models.

| Model | F1 | AUC | PR_AUC | MCC | Triad Overlap | Permutation Z | Score | Negative Overlap |
| --- | --- | --- | --- | --- | --- | --- | --- | --- |
| M1: ESM-2 + XGBoost | 0,91 | 0,99 | 0,90 | 0,90 | 0,5 | 1,76 | 1,83 | 1 |
| M4: ProtT5 + RF | 0,83 | 0,98 | 0,85 | 0,81 | 1,5 | 5,63 | 3,17 | 1 |
| M2: ESM-2 + RF | 0,79 | 0,98 | 0,87 | 0,76 | 0,5 | 4,05 | 3,33 | 0 |
| M7: Graph (chain edges) | 0,89 | 0,98 | 0,85 | 0,87 | 0 | -0,27 | 3,67 | 1 |
| M3: ProtT5 + XGBoost | 0,77 | 0,98 | 0,83 | 0,75 | 1 | 7,48 | 4,17 | 0 |
| M8: Graph (contact-map edges) | 0,84 | 0,95 | 0,64 | 0,81 | 0 | 0,48 | 5,17 | 2 |
| M5: AAC/CTD + XGBoost | 0,78 | 0,97 | 0,77 | 0,73 | 0 | 0,32 | 6,00 | 0 |
| M6: AAC/CTD + RF | 0,73 | 0,95 | 0,72 | 0,68 | 0 | -0,43 | 7,33 | 0 |

Notes: F1 = F1 score; AUC = Area Under the ROC Curve; PR\_AUC = Area Under the Precision-Recall Curve; MCC = Matthews Correlation Coefficient; Triad Overlap = mean of the catalytic-triad residues correctly recovered within a model's top-15 ranked positions across IsPETase and LCC; Permutation Z = permutation-test z-score for the combined triple-mutation effect on the catalytic triad relative to random residue triplets across IsPETase and LCC; Negative Overlap = number (out of 3) of catalytic residues recovered from the CalB negative-control reference; Score = composite rank, calculated as the mean of each model's individual ranks across F1, AUC, PR-AUC, MCC, Triad Overlap, and Permutation Z (lower Score = better overall rank); Negative Overlap is excluded from the Score calculation. Models are ordered by ascending Score.

## 4 Discussion

The ablation comparison indicates that a well-tuned combination of a pretrained protein language model embedding and a gradient-boosted classifier (M1: ESM-2 + XGBoost) remains the strongest overall predictor of PETase activity on this curated benchmark, outperforming both classical handcrafted features and graph-based architectures across F1, AUC, and MCC, and doing so with the tightest confusion-matrix error profile of all eight models (3 false positives, 1 false negative out of 139 benchmark sequences; Table S4). Notably, the narrow spread of ROC-AUC across all eight models (0.951–0.987) contrasts sharply with the much wider spread in average precision (0.641–0.905; Figure S4), underscoring that ROC-AUC alone would have obscured meaningful differences in model quality on this imbalanced benchmark (22 positive vs. 117 negative sequences) and reinforcing the choice to prioritize F1, PR-AUC, and MCC as the primary benchmark criteria (Imani, Beikmohammadi, and Arabnia 2025, Imani *et al*. 2026). Within the graph family, the higher F1 of M8 (contact-map edges, F1 = 0.840) over M7 (sequence-chain edges, F1 = 0.824) constitutes relatively clean evidence that explicit 3D structural contacts contribute predictive signal beyond sequence adjacency alone. However, this comes at a cost, as M8 also showed the lowest PR-AUC (0.64) and the widest confidence interval among the two graph variants (Table S3), suggesting that structural contact information improves discrimination on some subset of cases while simultaneously making the model’s confidence estimates less precise (Engelhard *et al*. 2025).

The feature ranking and directional effects observed in the SHAP analysis provide biophysically meaningful insights into the sequence determinants of PETase activity. For the AAC/CTD-based models, SHAP directly identified proline composition (AAC_P) as the dominant feature in both M5 (mean |SHAP| = 0.894) and M6, followed consistently by solvent accessibility, polarity, and secondary-structure composition. Proline overrepresentation across the entire sequence may impose excessive rigidity that compromises the conformational flexibility required for catalytic function. This is consistent with the trade-off between stability and activity often observed in PETase engineering, where proline substitutions near the active site can reduce activity despite improving thermostability (Prajapati *et al*. 2007, Umumararungu *et al*. 2024). The negative contributions of solvent accessibility and polarity (SolventAccessibility_T12 and Polarity_T13) are consistent with the hydrophobic nature of the PET substrate (Carr, Clarke, and Dobson 2020). Conversely, the positive contribution of leucine composition (AAC_L) and hydrophobicity descriptors supports the importance of hydrophobic residues in PETase activity (Karaoli *et al*. 2025). The positive contribution of SecondaryStructure_C3 further indicates that specific secondary-structure elements, likely β-strands or loops with defined geometry, are important for maintaining the canonical α/β-hydrolase fold that positions the catalytic triad (Ser160-Asp206-His237) optimally for catalysis (Kim *et al*. 2025).

For the embedding-based models, where individual ESM-2 and ProtT5 dimensions carry no inherent biophysical label, the cross-representation validation against 167 AAC/CTD descriptors provides an important bridge. A substantial fraction of each model’s top SHAP-important dimensions remained significantly correlated with named biophysical properties, suggesting that ESM-2 and, to a lesser extent, ProtT5 embeddings implicitly re-encode a meaningful share of the same biophysical information that AAC/CTD features encode explicitly. However, the graph-based models stood apart from this pattern. Despite comparable raw correlation counts, label-controlling produced the largest shrinkage in both graph variants (M7: 0.113; M8: 0.117) and zero dimensions meeting the strong-match criterion, indicating that the apparent AAC/CTD correspondence for M7 and M8 is disproportionately explained by both representations independently tracking the class label rather than reflecting a shared biophysical encoding. This is consistent with Kamal et al. (2025) which says that GNNs have demonstrated strong performance in prediction, but their interpretability remains limited.

The biological validation results substantially complicate a benchmark-only interpretation of model quality. Despite M1’s superior benchmark performance, it recovered only one catalytic-triad residue for IsPETase and one for LCC, corresponding to a modest mean overlap (0.5/3) and mean permutation z-score (1.76) across the two positive controls. Meanwhile M7, despite achieving the second-highest F1 score among all eight models, showed zero catalytic-triad overlap for both IsPETase and LCC and a negative or near-zero permutation z-score (IsPETase z = −0.25; LCC z = −0.29), indicating triad-region importance indistinguishable from chance. This dissociation between predictive accuracy and mechanistic grounding demonstrates that a model can achieve strong classification performance while relying substantially on signal unrelated to the experimentally confirmed active site. In the context of PETase, mutational effects have been observed to arise from residues distal to the active site, further illustrating that high predictive performance does not guarantee that a model’s decisions reflect experimentally confirmed catalytic determinants (Vongsouthi *et al*. 2025).

Conversely, M4 (ProtT5 + Random Forest) showed the strongest and most consistent triad recovery among all eight models. M4 recovering 1/3 residues for IsPETase (z = 4.36, p = 0.01) and 2/3 for LCC (z = 6.89, p < 0.01) despite a middling benchmark rank, suggesting that ProtT5 embeddings paired with Random Forest may encode positional or residue-level information more aligned with the true catalytic mechanism than the higher-scoring ESM-2 + XGBoost combination. The CalB negative-control results add a further layer of nuance. M8’s combination of zero true-triad overlap (for both IsPETase and LCC) with the highest off-target overlap and by far the largest z-score observed across the entire analysis (2/3 CalB residues recovered, z = 15.74, p < 0.01) suggests that the top-ranked positions identified by this model’s mutagenesis scan are disproportionately driven by generic α/β-hydrolase active-site geometry as a feature shared across many serine hydrolases regardless of PET-degrading capability rather than by PETase-specific mechanism (Ozhelvaci and Steczkiewicz 2025). Interestingly, several other models (M1, M4, M7) also showed statistically significant, if smaller, sensitivity to the CalB triad (z = 2.08– 4.74), indicating that some degree of generic active-site sensitivity is not unique to the graph architecture.

The composite ranking, by design, formalizes this tension between predictive performance and biological plausibility. M1’s first-place overall rank is driven almost entirely by its benchmark dominance, while its comparatively weak biological-validity standing (rank 3rd on triad overlap, 4th on permutation z among the eight models) is diluted by unweighted averaging across all six ranking criteria. For those who prioritizing mechanistic interpretability over raw accuracy alone might reasonably favor M4 instead, which ranked first on both biological-validity criteria despite only a middling benchmark rank. This highlights an important limitation of any single composite score: equal weighting across six criteria is a modeling choice rather than a ground truth, and different downstream applications may warrant different weightings. We therefore present the composite score primarily as a structured summary for cross-model comparison rather than as a definitive verdict on which single model should be adopted for all use cases.

Several limitations should be considered when interpreting these results. First, the benchmark set, while carefully curated via CD-HIT-2D filtering and near-duplicate removal to avoid data leakage (Figure S1-S2), remains modest in size (n = 139, with only 22 positive examples), which limits the statistical power of both the benchmark metrics and the mutagenesis-based permutation tests, as reflected in the comparatively wide bootstrap confidence intervals reported for several models (e.g., M2 and M3 F1 intervals spanning roughly 0.64–0.89; Table S3). Second, the mutagenesis analysis was restricted to three reference enzymes with experimentally confirmed catalytic residues. While IsPETase and LCC represent two independently characterized PET-hydrolyzing enzymes and CalB provides a mechanistically matched negative control, broader validation across additional reference structures would strengthen confidence in the generalizability of the biological-validity rankings. Third, the cross-representation correlation analysis is explicitly correlational rather than causal. A significant partial correlation between an embedding dimension and a named biophysical descriptor indicates shared statistical structure, not that the model uses that property in a mechanistic sense, and this limitation is compounded for the graph-based models, where the near-total loss of strong correspondence after label-controlling suggests that even this correlational evidence should be interpreted with particular caution for M7 and M8.

## 5 Conclusion

This study presents InterPET, a curated benchmark for PETase activity prediction that jointly evaluates sequence representation, classifier architecture, structural information, and biological validity, an integration rarely achieved in prior ML-based PETase discovery efforts. Across eight model configurations, ESM-2 embeddings paired with XGBoost achieved the strongest benchmark performance (F1 = 0.91, AUC = 0.99, MCC = 0.90). Meanwhile, the structure-aware graph models using 3D contact-map edges modestly outperformed their sequence-only counterparts, confirming that explicit structural information contributes predictive signal beyond sequence adjacency alone.

Critically, biological validation through SHAP-based cross-representation analysis and in-silico alanine-scanning mutagenesis revealed that benchmark accuracy alone is an incomplete proxy for mechanistic trustworthiness. This shown by the top-performing model by benchmark metrics recovered only half of the known catalytic-triad residues, while a model with only middling benchmark rank (ProtT5 + Random Forest) most consistently recovered catalytic-triad positions and showed no comparable sensitivity to an unrelated hydrolase’s active site. This dissociation, formalized through a composite ranking framework, demonstrates that predictive performance and biological plausibility are complementary but distinct axes of model quality, and that relying on a single leaderboard metric risks selecting models that classify accurately for the wrong reasons. These results argue that future PETase and, more broadly, enzyme-function prediction pipelines should report benchmark performance, interpretability, and biological-validity metrics side by side rather than optimizing for accuracy alone.

## Acknowledgements

None declared.

## Supplementary data

Supplementary data are available at *Bioinformatics Advances* online.

## Conflict of interest

The authors declare no competing interests.

## Funding

The authors received no specific funding for this work

## Data availability

The final dataset and code underlying this article are available in *Github* at https://github.com/indiraprakoso/interpet/

## Notes

https://github.com/indiraprakoso/interpet/

### Competing Interest Statement

The authors have declared no competing interest.

