## Supplementary Materials for "InterPET: A Curated Benchmark of Sequence Embeddings and Graph Architectures with Interpretability and Biological Validation for PETase Activity Prediction"

**Table S1.** Hyperparameter tuning selected for each of the eight ablation models. For each model (#1–#8, corresponding to M1–M8), the selected hyperparameters from the tuning search and the classification decision threshold used to convert predicted probabilities into binary labels are reported.

| Model | Selected Hyperparameters | Threshold |
|---|---|---|
| #1 | max_depth = 6, learning_rate = 0.05, n_estimators = 300, subsample = 0.8, colsample_bytree = 0.8 | 0.56 |
| #2 | n_estimators = 300, max_depth = None, min_samples_leaf = 3, max_features = "sqrt" | 0.31 |
| #3 | max_depth = 4, learning_rate = 0.05, n_estimators = 300, subsample = 0.8, colsample_bytree = 0.8 | 0.07 |
| #4 | n_estimators = 300, max_depth = 10, min_samples_leaf = 2, max_features = "sqrt" | 0.29 |
| #5 | max_depth = 6, learning_rate = 0.05, n_estimators = 300, subsample = 0.8, colsample_bytree = 0.8 | 0.57 |
| #6 | n_estimators = 300, max_depth = 10, min_samples_leaf = 2, max_features = "sqrt" | 0.51 |
| #7 | Hidden_channels = 128, num_layers = 3, dropout = 0.5 | 0.78 |
| #8 | hidden_channels = 64, num_layers = 3, dropout = 0.3 | 0.58 |

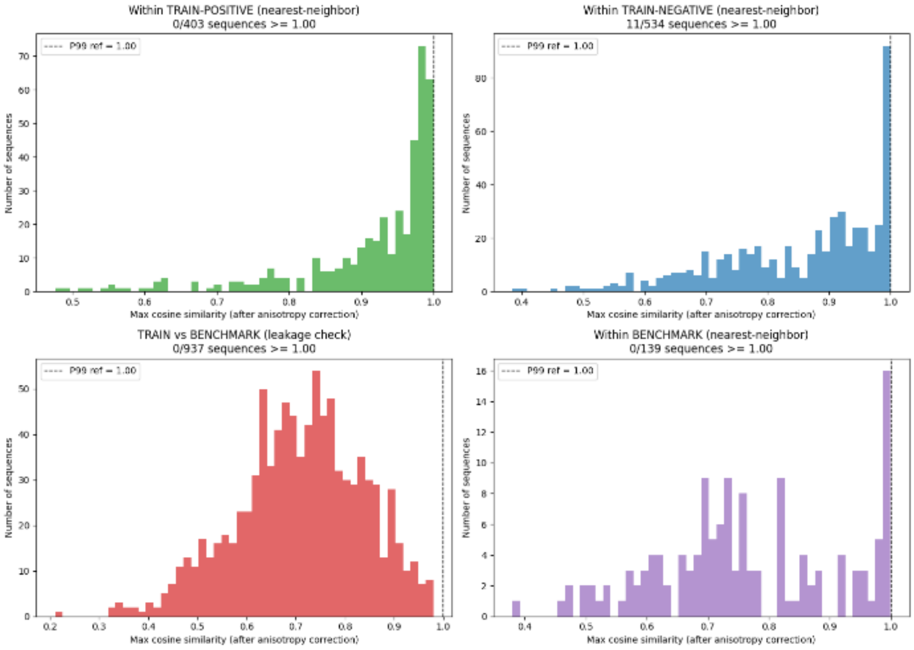


**Figure S1.** Distribution of maximum cosine similarity between ESM-2 embeddings after anisotropy correction: (Upper left) within training positive sequences, (Upper right) within training negative sequences, (Lower left) training versus benchmark sequences, and (Lower right) within benchmark sequences. The vertical dashed line indicates the threshold for potential near-duplicates (P99 = 1.000).

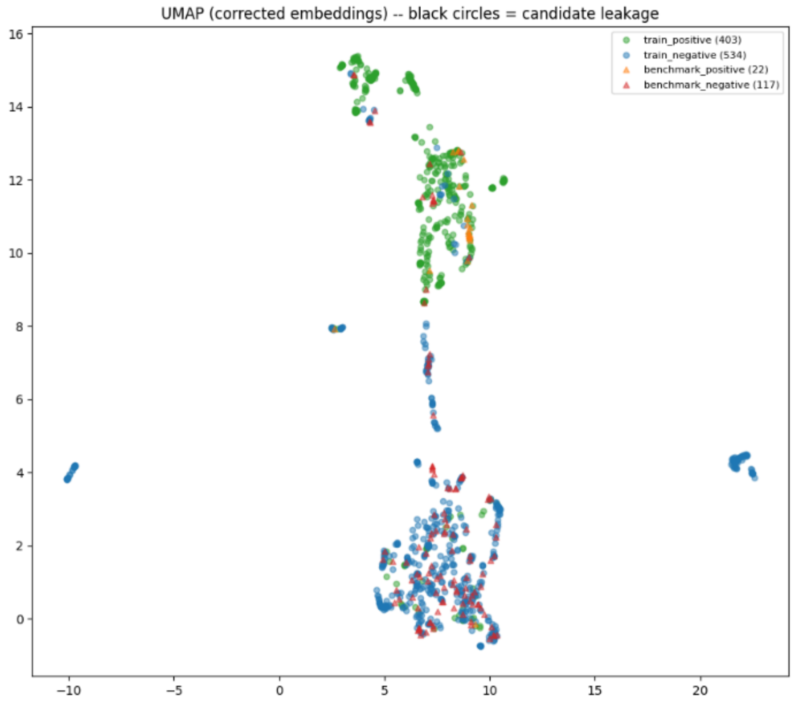


**Figure S2**. UMAP projection of corrected embeddings for training and benchmark samples. Green and blue points denote positive (n = 403) and negative (n = 534) training examples, respectively; orange and red triangles denote positive (n = 22) and negative (n = 117) benchmark examples. Black-outlined markers indicate candidate leakage points.

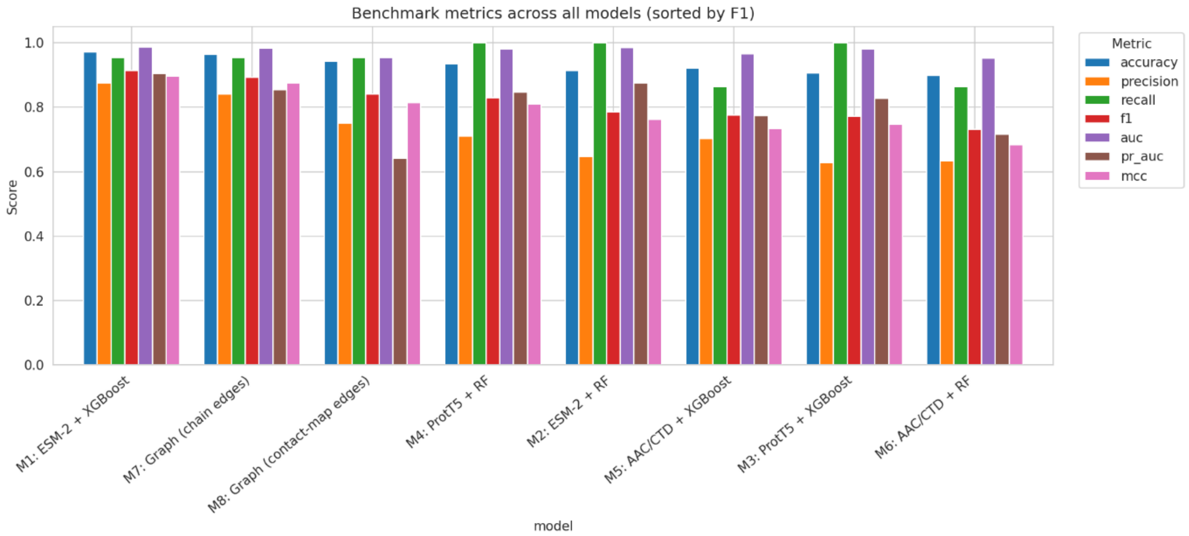


**Figure S3.** Benchmark metrics across all eight ablation models (M1–M8), sorted by F1 score. Bars show accuracy, precision, recall, F1, AUC, PR-AUC, and MCC evaluated on the independent benchmark set.

**Table S2.** Benchmark performance metrics (accuracy, precision, recall, F1, AUC, PR-AUC, MCC) for all eight ablation models (M1–M8) evaluated on the independent benchmark set, ranked by F1 score.

| model | accuracy | precision | recall | f1 | auc | pr_auc | mcc |
|---|---|---|---|---|---|---|---|
| M1: ESM-2 + XGBoost | 0.97 | 0.88 | 0.95 | 0.91 | 0.99 | 0.90 | 0.90 |
| M7: Graph (chain edges) | 0.96 | 0.84 | 0.95 | 0.89 | 0.98 | 0.85 | 0.87 |
| M8: Graph (contact-map edges) | 0.94 | 0.75 | 0.95 | 0.84 | 0.95 | 0.64 | 0.81 |
| M4: ProtT5 + RF | 0.94 | 0.71 | 1.00 | 0.83 | 0.98 | 0.85 | 0.81 |
| M2: ESM-2 + RF | 0.91 | 0.65 | 1.00 | 0.79 | 0.98 | 0.87 | 0.76 |
| M5: AAC/CTD + XGBoost | 0.92 | 0.70 | 0.86 | 0.78 | 0.97 | 0.77 | 0.73 |
| M3: ProtT5 + XGBoost | 0.91 | 0.63 | 1.00 | 0.77 | 0.98 | 0.83 | 0.75 |
| M6: AAC/CTD + RF | 0.90 | 0.63 | 0.86 | 0.73 | 0.95 | 0.72 | 0.68 |

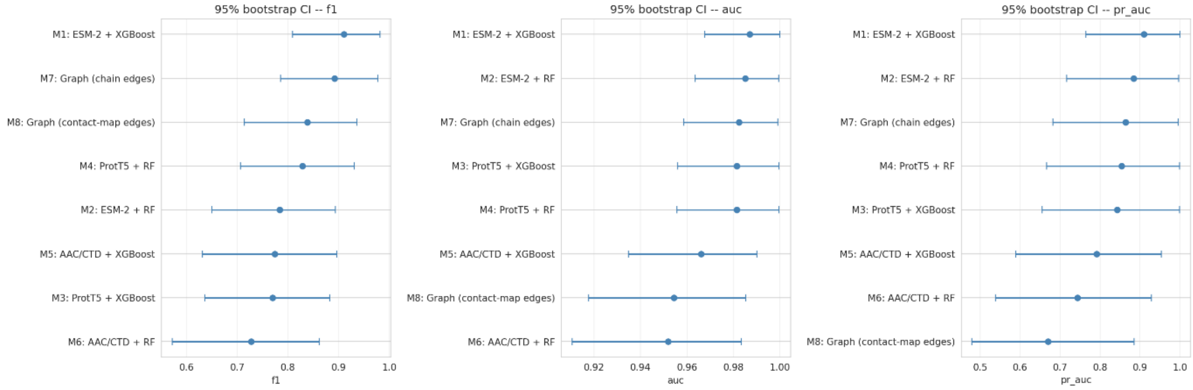


**Figure S4.** Bootstrap 95% confidence intervals for F1 (left), AUC (center), and PR-AUC (right) across all eight ablation models (M1–M8), estimated by resampling the benchmark set. Points indicate the mean value; error bars indicate the 95% confidence interval.

**Table S3.** Bootstrap 95% confidence intervals (mean, lower and upper bounds) for F1, precision, recall, AUC, and PR-AUC for each of the eight ablation models (M1–M8) on the benchmark set.

| Model | Metric | Mean | CI_low | CI_high |
|---|---|---|---|---|
| M1: ESM-2 + XGBoost | f1 | 0.91 | 0.81 | 0.98 |
| M1: ESM-2 + XGBoost | precision | 0.87 | 0.73 | 1.00 |
| M1: ESM-2 + XGBoost | recall | 0.95 | 0.83 | 1.00 |
| M1: ESM-2 + XGBoost | auc | 0.99 | 0.97 | 1.00 |
| M1: ESM-2 + XGBoost | pr_auc | 0.91 | 0.76 | 1.00 |
| M2: ESM-2 + RF | f1 | 0.78 | 0.65 | 0.89 |
| M2: ESM-2 + RF | precision | 0.65 | 0.48 | 0.81 |
| M2: ESM-2 + RF | recall | 1.00 | 1.00 | 1.00 |
| M2: ESM-2 + RF | auc | 0.98 | 0.96 | 1.00 |
| M2: ESM-2 + RF | pr_auc | 0.88 | 0.72 | 1.00 |
| M3: ProtT5 + XGBoost | f1 | 0.77 | 0.64 | 0.88 |
| M3: ProtT5 + XGBoost | precision | 0.63 | 0.47 | 0.79 |
| M3: ProtT5 + XGBoost | recall | 1.00 | 1.00 | 1.00 |
| M3: ProtT5 + XGBoost | auc | 0.98 | 0.96 | 1.00 |
| M3: ProtT5 + XGBoost | pr_auc | 0.84 | 0.66 | 1.00 |
| M4: ProtT5 + RF | f1 | 0.83 | 0.71 | 0.93 |
| M4: ProtT5 + RF | precision | 0.71 | 0.55 | 0.87 |
| M4: ProtT5 + RF | recall | 1.00 | 1.00 | 1.00 |
| M4: ProtT5 + RF | auc | 0.98 | 0.96 | 1.00 |
| M4: ProtT5 + RF | pr_auc | 0.85 | 0.67 | 1.00 |
| M5: AAC/CTD + XGBoost | f1 | 0.77 | 0.63 | 0.90 |
| M5: AAC/CTD + XGBoost | precision | 0.71 | 0.53 | 0.88 |
| M5: AAC/CTD + XGBoost | recall | 0.87 | 0.71 | 1.00 |
| M5: AAC/CTD + XGBoost | auc | 0.97 | 0.93 | 0.99 |
| M5: AAC/CTD + XGBoost | pr_auc | 0.79 | 0.59 | 0.95 |
| M6: AAC/CTD + RF | f1 | 0.73 | 0.57 | 0.86 |
| M6: AAC/CTD + RF | precision | 0.64 | 0.46 | 0.82 |
| M6: AAC/CTD + RF | recall | 0.86 | 0.70 | 1.00 |
| M6: AAC/CTD + RF | auc | 0.95 | 0.91 | 0.98 |
| M6: AAC/CTD + RF | pr_auc | 0.74 | 0.54 | 0.93 |
| M7: Graph (chain edges) | f1 | 0.89 | 0.79 | 0.98 |
| M7: Graph (chain edges) | precision | 0.84 | 0.69 | 0.96 |
| M7: Graph (chain edges) | recall | 0.95 | 0.83 | 1.00 |
| M7: Graph (chain edges) | auc | 0.98 | 0.96 | 1.00 |
| M7: Graph (chain edges) | pr_auc | 0.86 | 0.68 | 1.00 |
| M8: Graph (contact-map edges) | f1 | 0.84 | 0.71 | 0.94 |
| M8: Graph (contact-map edges) | precision | 0.75 | 0.59 | 0.90 |
| M8: Graph (contact-map edges) | recall | 0.95 | 0.83 | 1.00 |
| M8: Graph (contact-map edges) | auc | 0.95 | 0.92 | 0.99 |
| M8: Graph (contact-map edges) | pr_auc | 0.67 | 0.48 | 0.89 |

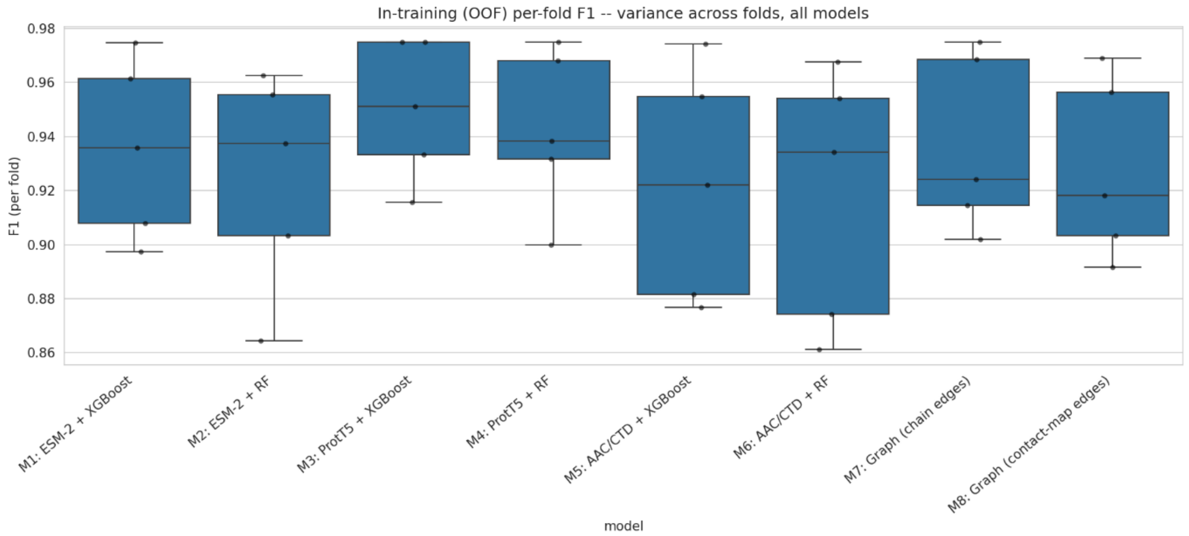


**Figure S5.** Cross-validation fold metrics. Distribution of out-of-fold (OOF) F1 scores across cross-validation folds for each of the eight ablation models (M1–M8), illustrating the variance in training performance across folds.

**Table S4.** Mean and standard deviation of out-of-fold (OOF) F1 scores across cross-validation folds for each of the eight ablation models (M1–M8).

| **model** | **mean** | **std** |
|---|---|---|
| M1: ESM-2 + XGBoost | 0.935490 | 0.033258 |
| M2: ESM-2 + RF | 0.924631 | 0.040679 |
| M3: ProtT5 + XGBoost | 0.950043 | 0.026021 |
| M4: ProtT5 + RF | 0.942620 | 0.030230 |
| M5: AAC/CTD + XGBoost | 0.921913 | 0.043315 |
| M6: AAC/CTD + RF | 0.918297 | 0.047979 |
| M7: Graph (chain edges) | 0.936840 | 0.032921 |
| M8: Graph (contact-map edges) | 0.927730 | 0.033596 |

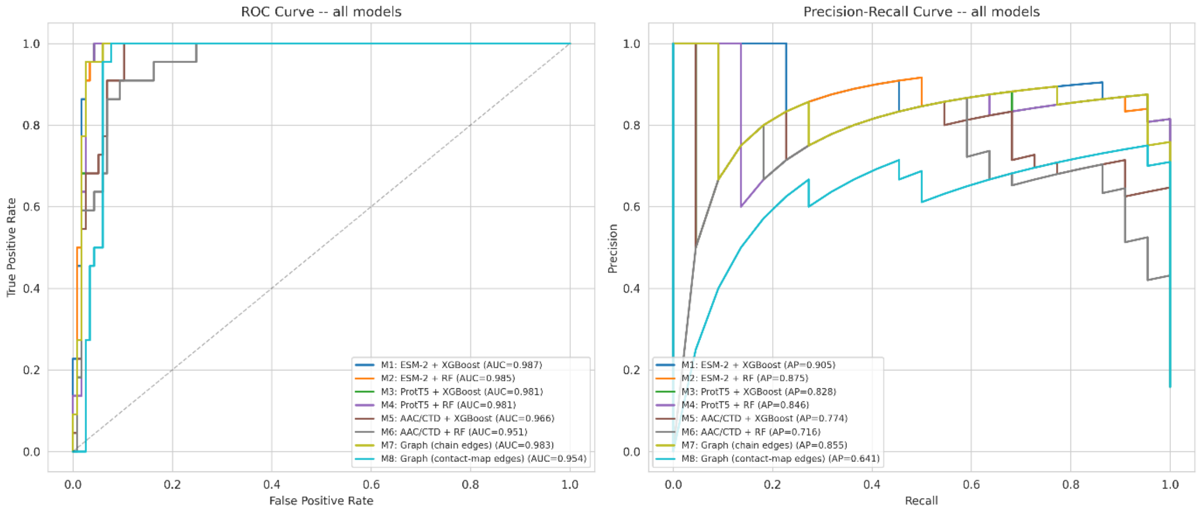


**Figure S6.** ROC (left) and Precision–Recall (right) curves for all eight ablation models (M1–M8) evaluated on the benchmark set. The area under the ROC curve (AUC) and average precision (AP) are reported in the legend for each model.

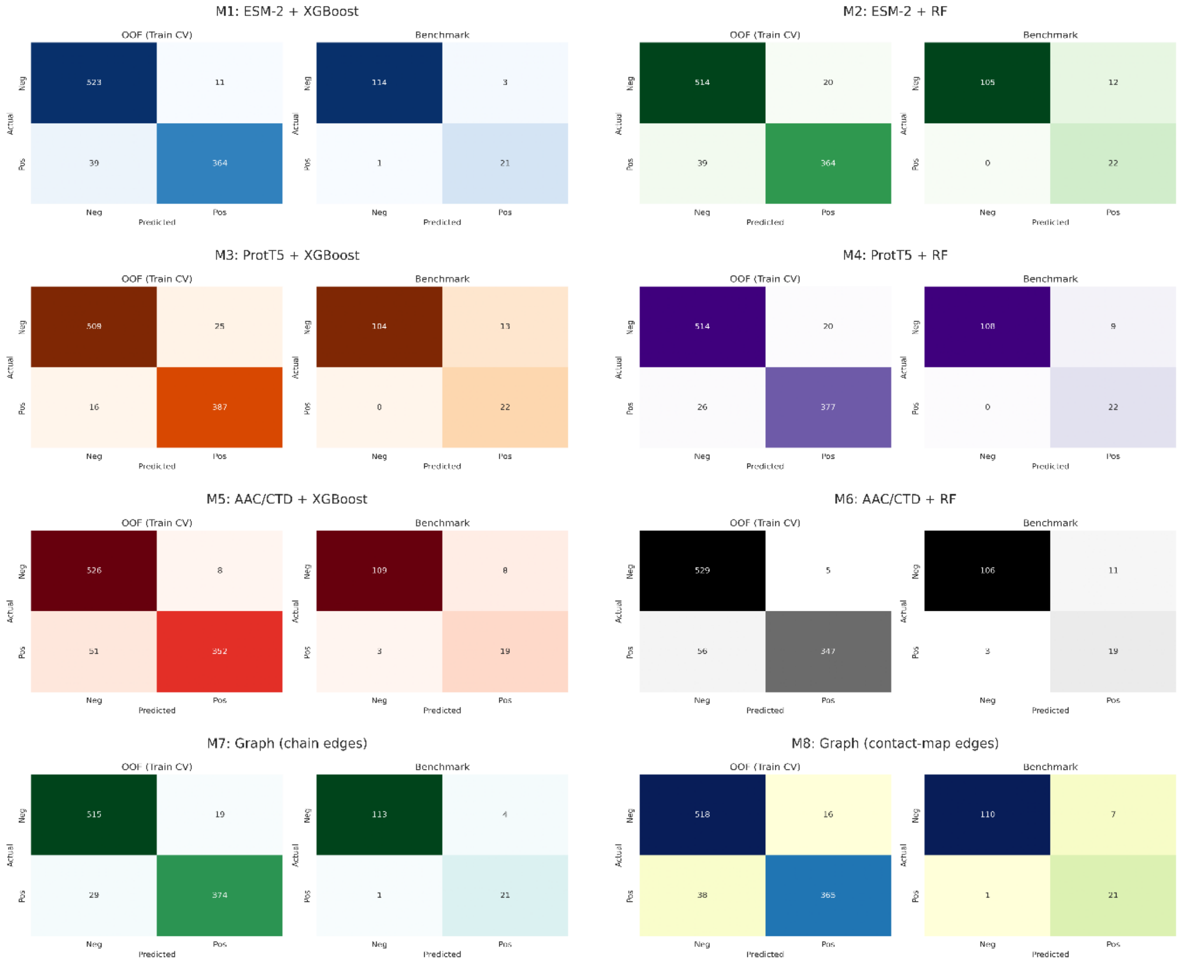


**Figure S7.** Confusion matrices comparing out-of-fold (OOF) training predictions and independent benchmark predictions for each of the eight ablation models (M1–M8). True negative, false positive, false negative, and true positive counts are shown for both the training (OOF) and benchmark sets.

**Table S5.** Confusion matrix comparison (true positives, false positives, false negatives, true negatives) on the independent benchmark set for each of the eight ablation models (M1–M8).

| model | TP | FP | FN | TN |
|---|---|---|---|---|
| M1: ESM-2 + XGBoost | 21 | 3 | 1 | 114 |
| M2: ESM-2 + RF | 22 | 12 | 0 | 105 |
| M3: ProtT5 + XGBoost | 22 | 13 | 0 | 104 |
| M4: ProtT5 + RF | 22 | 9 | 0 | 108 |
| M5: AAC/CTD + XGBoost | 19 | 8 | 3 | 109 |
| M6: AAC/CTD + RF | 19 | 11 | 3 | 106 |
| M7: Graph (chain edges) | 21 | 4 | 1 | 113 |
| M8: Graph (contact-map edges) | 21 | 7 | 1 | 110 |

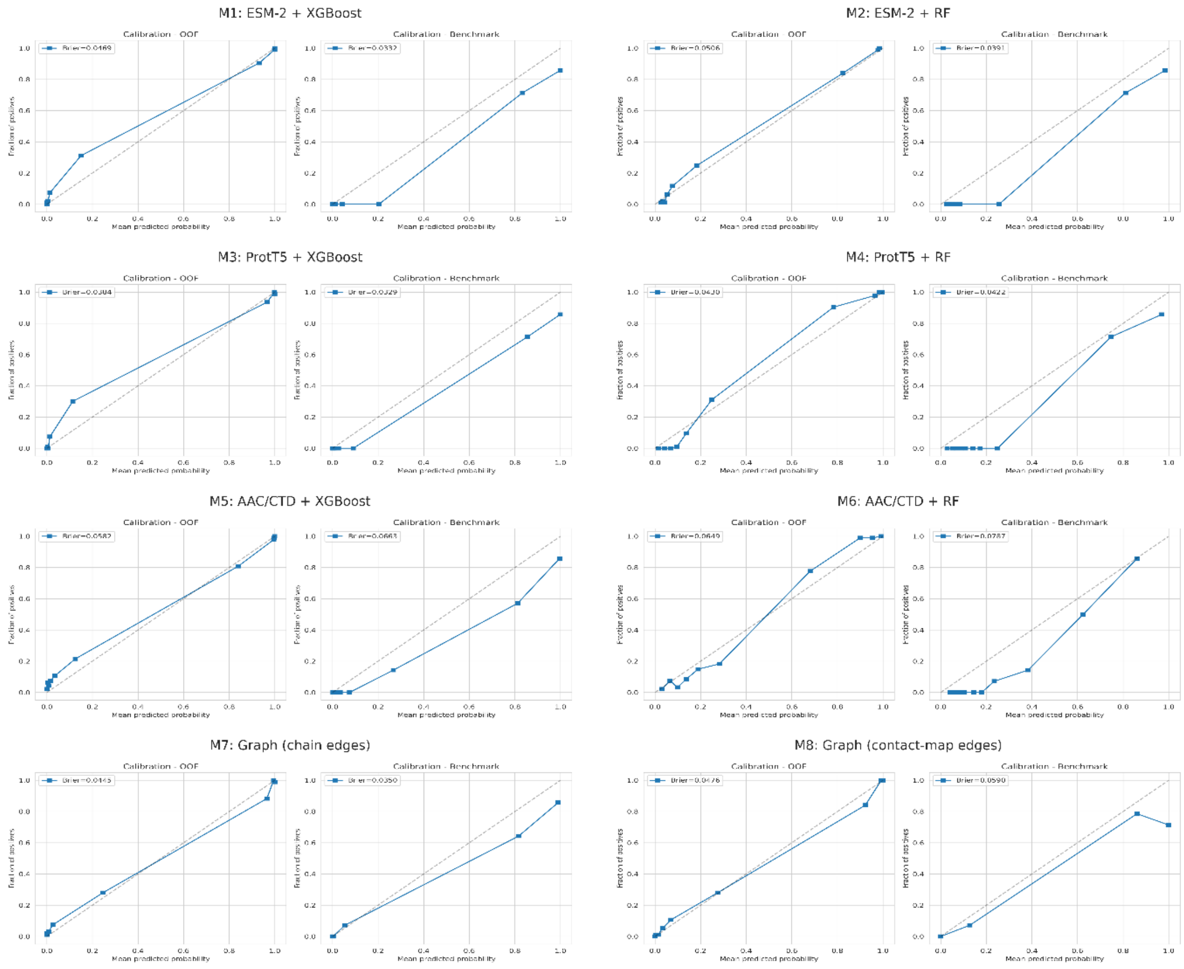


**Figure S8.** Calibration curves and Brier scores for out-of-fold (OOF) training predictions and independent benchmark predictions for each of the eight ablation models (M1–M8). The dashed diagonal line indicates perfect calibration; the Brier score quantifies the mean squared difference between predicted probabilities and observed outcomes.

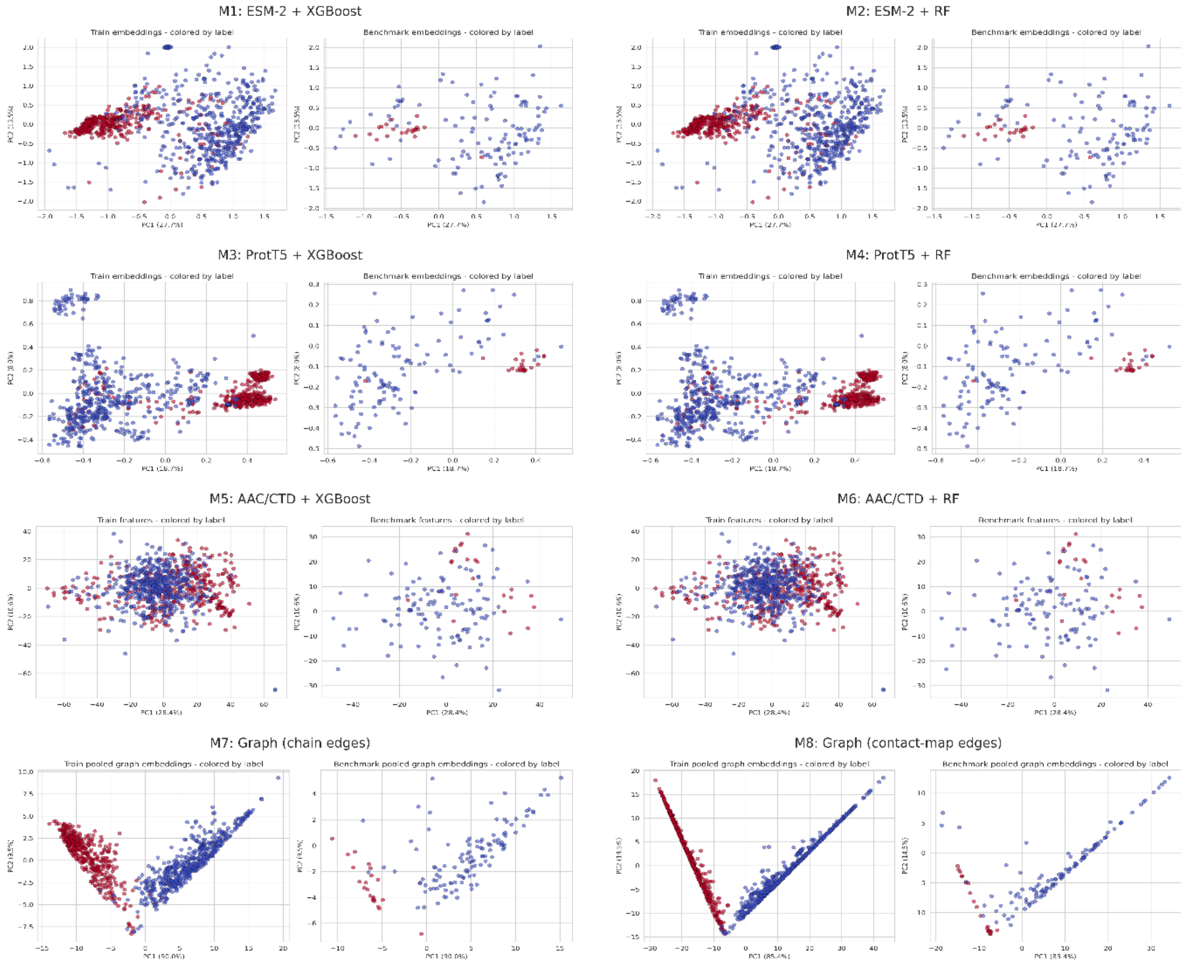


**Figure S9.** Principal component analysis (PCA) of the input representation. Each of the eight ablation models (M1-M8), comparing the training set (left panel) and the benchmark set (right panel), colored by class label (red = positive, blue = negative). Percentages on each axis indicate the proportion of variance explained by that principal component.

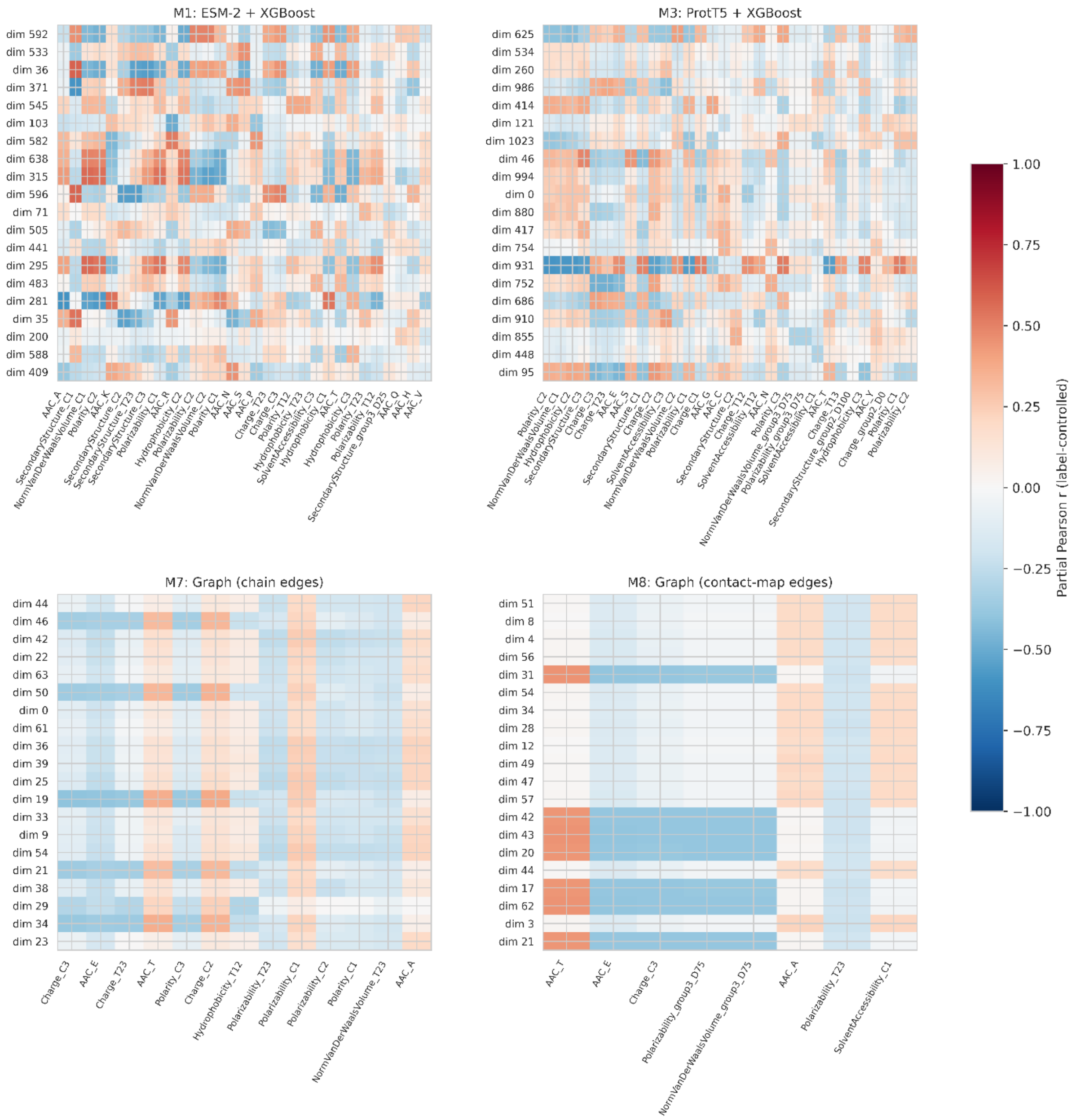


**Figure S10.** Cross-representation validation of top SHAP-important embedding dimensions against known biophysical descriptors, shown as label-controlled partial Pearson correlation heatmaps for ESM-2 (upper left), ProtT5 (Upper right), Graph with chain-edges (Lower left), Graph with contact-map edges (Lower right). Rows represent each model's top-ranked embedding dimensions (by mean absolute SHAP value); columns represent AAC/CTD-derived biophysical descriptors (amino-acid composition, charge, polarity, hydrophobicity, polarizability, secondary-structure propensity, solvent accessibility, and normalized van der Waals volume). Color indicates the partial Pearson correlation coefficient between each embedding dimension and each descriptor. Red indicates positive correlation, blue indicates negative correlation. This analysis provides correlational, not mechanistic, evidence for the biophysical properties putatively captured by each model's top SHAP-important dimensions.

**Table S6.** In-silico alanine-scanning mutagenesis results for each ablation model (M1–M8) evaluated against three reference PETase structures (CalB, IsPETase, LCC). Columns report the model's predicted probability for the reference sequence (base_proba), the number of catalytic-triad residues among the top 15 SHAP-ranked positions (top15_overlap_with_triad), the best-ranked catalytic-triad residue (best_triad_rank), the combined change in predicted probability upon triple alanine substitution of the catalytic triad (combined_triple_mutation_delta), and the permutation-based p-value and z-score assessing whether the triad's importance exceeds that of randomly selected residue sets.

| model | reference | base_proba | top15_overlap_with_triad | best_triad_rank | combined_triple_mutation_delta | permutation_p_value | permutation_z_score |
|---|---|---|---|---|---|---|---|
| M8: Graph (contact-map edges) | CalB | 0.99 | 2 | 1 | 0.15 | 0.00 | 15.74 |
| M1: ESM-2 + XGBoost | CalB | 0.96 | 1 | 12 | 0.17 | 0.02 | 2.81 |
| M4: ProtT5 + RF | CalB | 0.66 | 1 | 3 | 0.18 | 0.00 | 4.74 |
| M7: Graph (chain edges) | CalB | 0.23 | 1 | 2 | 0.19 | 0.00 | 2.08 |
| M2: ESM-2 + RF | CalB | 0.57 | 0 | 16 | 0.15 | 0.02 | 2.72 |
| M3: ProtT5 + XGBoost | CalB | 0.96 | 0 | 18 | 0.07 | 0.43 | 0.03 |
| M5: AAC/CTD + XGBoost | CalB | 0.99 | 0 | 105 | 0.01 | 0.52 | -0.32 |
| M6: AAC/CTD + RF | CalB | 0.84 | 0 | 77 | 0.04 | 0.41 | 0.20 |
| M3: ProtT5 + XGBoost | IsPETase | 1.00 | 1 | 2 | 0.00 | 0.00 | 7.17 |
| M4: ProtT5 + RF | IsPETase | 1.00 | 1 | 12 | 0.04 | 0.01 | 4.36 |
| M1: ESM-2 + XGBoost | IsPETase | 1.00 | 0 | 23 | 0.00 | 0.01 | 3.18 |
| M2: ESM-2 + RF | IsPETase | 0.98 | 0 | 170 | 0.01 | 0.04 | 1.37 |
| M5: AAC/CTD + XGBoost | IsPETase | 1.00 | 0 | 54 | 0.00 | 0.27 | 0.30 |
| M6: AAC/CTD + RF | IsPETase | 1.00 | 0 | 151 | 0.00 | 1.00 | -0.80 |
| M7: Graph (chain edges) | IsPETase | 0.99 | 0 | 50 | 0.00 | 0.63 | -0.25 |
| M8: Graph (contact-map edges) | IsPETase | 1.00 | 0 | 39 | 0.00 | 0.11 | 1.27 |
| M4: ProtT5 + RF | LCC | 1.00 | 2 | 1 | 0.03 | 0.00 | 6.89 |
| M1: ESM-2 + XGBoost | LCC | 1.00 | 1 | 11 | 0.00 | 0.15 | 0.33 |
| M2: ESM-2 + RF | LCC | 0.99 | 1 | 8 | 0.00 | 0.00 | 6.73 |
| M3: ProtT5 + XGBoost | LCC | 1.00 | 1 | 1 | 0.00 | 0.00 | 7.79 |
| M5: AAC/CTD + XGBoost | LCC | 1.00 | 0 | 57 | 0.00 | 0.29 | 0.35 |
| M6: AAC/CTD + RF | LCC | 1.00 | 0 | 23 | 0.01 | 0.42 | -0.06 |

| | | | | | | | |
|---|---|---|---|---|---|---|---|
| M7: Graph (chain edges) | LCC | 1.00 | 0 | 108 | 0.00 | 0.59 | -0.29 |
| M8: Graph (contact-map edges) | LCC | 1.00 | 0 | 145 | 0.00 | 0.46 | -0.30 |
